# Systemic Iron Signaling via OPT3 Influences Reductive Uptake and Coumarin Secretion

**DOI:** 10.64898/2026.01.02.696758

**Authors:** Chandan K Gautam

**Affiliations:** Institute of Plant and Microbial Biology, Academia Sinica, Taipei 11529, Taiwan

**Keywords:** OPT3, systemic signal, coumarins, crosstalk, Iron, nutrition

## Abstract

Iron (Fe) availability is frequently restricted in soils, particularly under alkaline conditions, resulting in Fe-deficiency chlorosis and reduced plant growth. In Strategy I plants, Fe acquisition relies on ferric chelate reductase (FCR) activity and the secretion of Fe-mobilizing compounds like coumarins; however, how these pathways are integrated through systemic Fe signaling remains poorly understood. In this study, coordination between reductive Fe uptake, coumarin biosynthesis, external pH, and systemic signaling was investigated in *Arabidopsis thaliana*. Physiological readouts, including chlorophyll content, root FCR activity, and coumarin fluorescence, were compared among wild-type Col-0, the systemic signaling mutant *opt3-2*, and the coumarin-deficient mutant *f6’h1-1* under Fe-sufficient, alkaline, and Fe-free conditions. Under alkaline Fe limitation, *opt3-2* sustained higher chlorophyll levels, persistent FCR activity, and strongly enhanced coumarin secretion, whereas *f6’h1-1* exhibited severe chlorosis accompanied by compensatory FCR induction. Time-resolved analyses showed that external pH restricts FCR responsiveness, while loss of OPT3-mediated systemic signaling profoundly alters the timing and amplitude of coumarin deployment. Co-cultivation assay further demonstrated that coumarins released by *opt3-2* partially alleviate Fe-deficiency symptoms in *f6’h1-1*. Together, these results identify systemic signaling as a key integrator of reductive Fe uptake and coumarin-mediated Fe mobilization, establishing a unified framework for Fe homeostasis.

---

Iron (Fe) is an essential micronutrient for plants, needed for photosynthesis, respiration, chlorophyll biosynthesis, and many Fe-dependent catalytic reactions. Its availability is often limited in soils, especially under alkaline conditions, where ferric (Fe^3+^) solubility is significantly reduced. This leads to Fe-deficiency chlorosis and reduced plant growth (**Clarkson, 1996**). Strategy I plants, which include dicots and non-graminaceous monocots, respond to this deficiency through a reduction-based Fe acquisition system. This system combines activation of plasma membrane-located ferric chelate reductase (FCR) to reduce Fe^3+^ to Fe^2+^, along with the secretion of Fe-mobilizing coumarins that chelate or reduce Fe in the rhizosphere (**Connolly et al., 2003; Schmid et al., 2014; Rajniak et al., 2018; Tsai et al., 2018; Gautam et al., 2021; Paffrath et al., 2024**). A recent finding by **Paffrath et al., 2024** has highlighted a central role for coumarin-dependent ferric Fe reduction in Strategy I Fe acquisition in Arabidopsis and suggested close crosstalk between FCR activity and coumarin biosynthesis, especially under alkaline conditions where classical reductive uptake alone is insufficient (**Tsai et al., 2018; Gautam et al., 2021; Paffrath et al., 2024**). At the heart of the Fe uptake pathway lies a fine-tuned systemic signaling pathway that relays whole-plant Fe status from shoot to roots. The phloem-localized transporter OPT3 (OLIGOPEPTIDE TRANSPORTER 3) is a key component of this systemic network (**Stacey et al., 2008)**. Loss of OPT3 results in misregulated Fe distribution and a constitutive Fe-deficiency response in roots, despite elevated shoot Fe levels, showing that systemic Fe signaling is fundamental for tuning local Fe uptake machinery (**Stacey et al., 2008**; **Mendoza-Cózatl et al., 2014; Zhai et al., 2014**). How these systemic signals coordinate the balance between FCR activity and coumarin biosynthesis, and how this balance is affected by external pH, remains poorly understood.

To investigate these questions, three well-established physiological proxies: chlorophyll content, FCR activity, and coumarin production, were used as robust, quantitative/qualitative readouts of Fe-deficiency responses in Arabidopsis. Using these accessible indicators, Fe-deficiency responses were examined in wild-type Col-0, *opt3-2* (Salk_021168C) mutant (described by **Stacey et al., 2008)** defective in shoot-to-root Fe-deficiency signaling, and in the *f6’h1-1* mutant (Salk_132418C) impaired in key coumarin biosynthesis protein Feruloyl-CoA 6′-Hydroxylase 1, F6’H1 (**Schmid et al., 2014**). Together, these well-characterized genetic backgrounds provide a sound framework to ask whether the coumarin pathway can be triggered in Fe-sufficient plants with distorted Fe distribution and systemic signaling (*opt3-2*), whether the loss of coumarin production in *f6’h1-1* can be compensated via improved reductive Fe uptake through FCR, how these responses relate quantitatively to chlorophyll maintenance under Fe-limiting and alkaline conditions, and whether the abundant coumarins produced by *opt3-2* are sufficient to rescue Fe-deficiency symptoms in the coumarin-deficient *f6’h1-1*. The three genotypes were first grown for 9 days (d) on **Estelle and Somerville (1987)** (ES/control) (40 µM Fe-EDTA, pH 5.5) medium and then transferred either back to ES or to non-available Fe (navFe/alkaline) (40 µM FeCl_3_, pH 7.0) and Fe-free (-Fe) (0 Fe, 120 µM Ferrozine, pH 5.5) nutrient agar plates, as described by **Gautam et al. (2021)**. Under control conditions, all genotypes showed comparable chlorophyll content, indicating similar chlorophyll status and absence of overt Fe deficiency symptoms (**Fig. 1A, B**), whereas *opt3-2* mutants displayed significantly higher FCR activity than Col-0 and *f6’h1-1*(**Fig. 1C**), consistent with previous reports that *opt3-2* roots maintain a constitutive Fe-deficiency signal and elevated FCR activity despite increased internal Fe levels (**Stacey et al., 2008; Mendoza-Cózatl et al., 2014**). This reiterates that disruption of systemic Fe signaling via OPT3 can uncouple tissue Fe status from root Fe-deficiency responses.

**Fig. 1.**
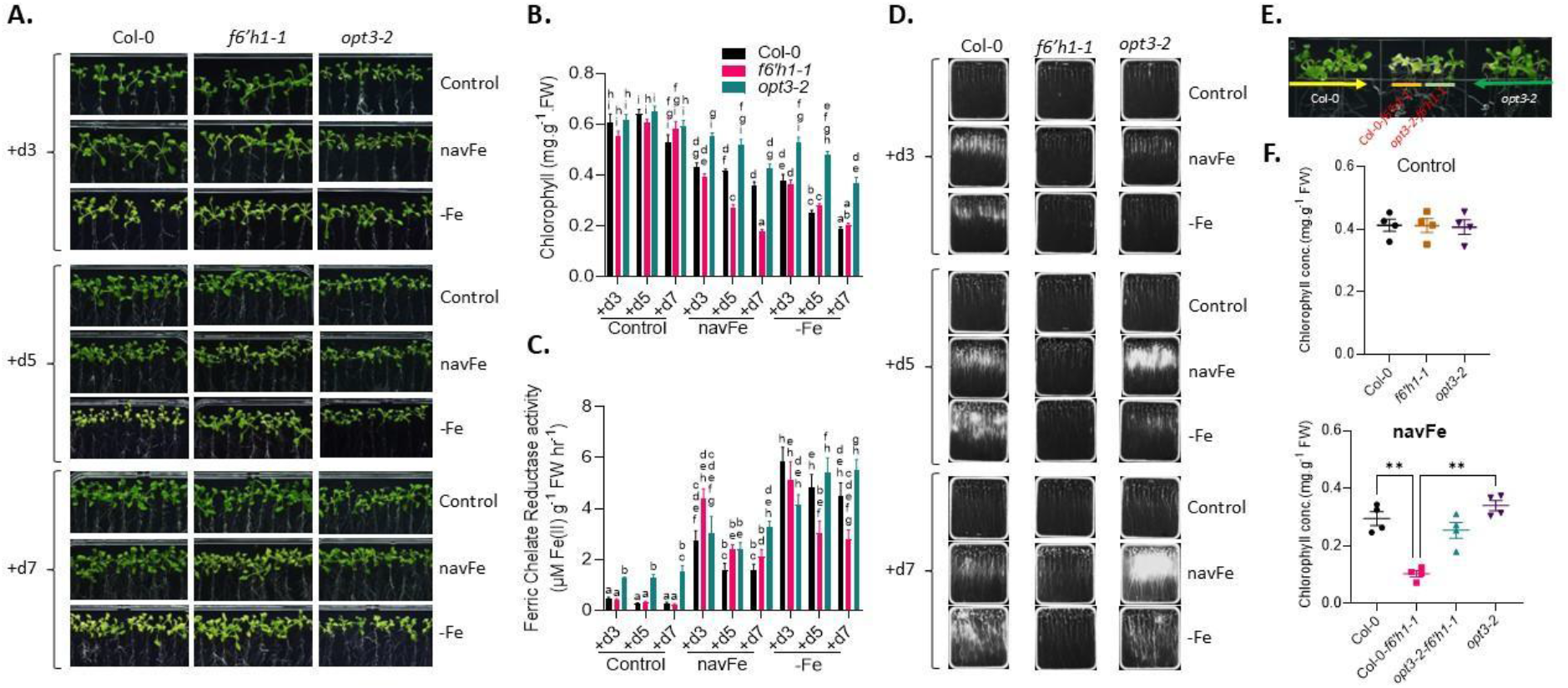
Growth phenotype, chlorophyll content, Ferric Chelate Reductase (FCR) activity, UV fluorescence imaging, and phenotype rescue. 9-d old seedlings (Col-0, *f6’h1-1, opt3-2*) grown on Control (ES) media were transferred to ES, alkaline (navFe), and Ferrozine(-Fe) containing nutrient media for 3day (d), 5d, and 7d. **A**. Pictures were taken under white light. Figures above are representative of three replicates. For measuring chlorophyll using the method developed by **Wellburn (1994)** (B), and FCR activity as described previously by **Grillet et al. (2014**) (**1C**), shoots and roots were harvested respectively at the end of the treatment. The data represent the average of three independent biological replicates, with two technical replicates per biological replicate. **D**. pictures showing fluorescent coumarins were captured under UV light 365 nm at the end of the experiment (as described by **Tsai et al., 2018**). Figures above are representative of three biological replicates. **E**. recue of coumarin-deficient mutant (*f6’h1-1*) on navFe media with Col-0 and *opt3-2* mutants. The *f6’h1-1* mutants were germinated and grown between Col-0 and *opt3-2* seedlings horizontally on navFe media for 14-d, following which chlorophyll was measured (**F**). The arrowheads in **(E)** denote the possible diffusion direction of coumarins. The figure and data are representative of four independent experiments. Control/ES (40 µM Fe-EDTA, pH 5.5); navFe/non-available Fe (40 µM FeCl_3_, pH 7.0), Ferrozine/-Fe (0 Fe, 120 µM Ferrozine, Fe free, pH 5.5), FW (Fresh weight). Distinct letters above the bars indicate statistically significant differences (P < 0.05) as determined by two-way ANOVA followed by Tukey’s HSD test. Asterisks above the plots indicate significant differences (*P < 0.05; **P < 0.01) as determined by one-way ANOVA followed by Tukey’s HSD test.

Upon transferring to navFe and -Fe media, genotype- and treatment-dependent patterns emerged. Under both navFe and -Fe conditions, *opt3-2* consistently exhibited the highest chlorophyll content, whereas *f6’h1-1* showed the lowest chlorophyll under navFe but similar levels as Col-0 under -Fe treatment, showing elevated Fe-deficiency tolerance in *opt3-2* and pronounced sensitivity of the coumarin-deficient *f6’h1-1* under alkaline conditions (**Fig. 1B**). At the mechanistic level, plants can increase Fe acquisition either by reducing non-available Fe^3+^ at the root surface via FCR and taking up Fe^2+^ through IRT-type transporters, or by secreting coumarins that mobilize Fe in the rhizosphere through chelation and redox chemistry (**Connolly et al., 2003; Schmid et al., 2014; Tsai et al., 2018; Spielmann et al., 2023; Paffrath et al., 2024**). FCR enzymatic activity of the freshly harvested roots, therefore, is a widely used proxy for the capacity to reduce Fe^3+^ and support uptake. At the same time, coumarin production can be conveniently monitored by imaging at 365 nm UV fluorescence, which reports the presence of fluorescent coumarins such as scopoletin in and around the roots (**Tsai et al., 2018; Gautam et al., 2021**). So, these two complementary approaches were used to capture the dynamics of reductive and chelator-based Fe acquisition in adaptation to changing Fe status and external pH.

Under navFe, all three genotypes showed a strong induction of FCR activity on day (d) 3, followed by a decline on d5 and d7, with *opt3-2* as a notable exception, maintaining a persistently similar FCR response throughout the time course (**Fig. 1C**). Under -Fe conditions, a similar early peak in FCR activity was observed for all genotypes on d3 of treatment; this response declined in *f6’h1-1* but remained elevated in *opt3-2* and, to a lesser extent, Col-0 (**Fig. 1C**). Across all genotypes, FCR activity was generally much higher under -Fe than under navFe, underscoring external pH as a key determinant of the magnitude and efficiency of FCR responses and reinforcing the notion that alkaline conditions restrict reductive Fe uptake more strongly than Fe-free but acidic conditions (**Fig. 1C**) **(Solti et al., 2014; Gautam et al., 2021; Paffrath et al., 2024**). In parallel, coumarin dynamics were visualized under 365 nm UV light. The navFe and -Fe treatment induced a fluorescent coumarin signal in both Col-0 and *opt3-2*, although the timing and intensity of the response differed between genotypes (**Fig. 1D**). In Col-0, coumarin fluorescence was already clearly detectable by d3 under navFe, indicating rapid deployment of coumarin-based Fe mobilization under alkaline Fe limitation (**Fig. 1D**). In contrast, in *opt3-2*, only a faint signal was visible at this time point. Consistent with previous findings, *f6’h1-1* did not show any fluorescent signals (**Schmid et al., 2014; Tsai et al., 2018; Gautam et al., 2021**). At d3 under navFe treatment, *opt3-2* did not yet display a dramatically higher FCR activity than Col-0, indicating that both coumarin secretion and FCR activity need a certain level and duration of Fe deficiency, to fully activate these responses, as supported by the pronounced increase in coumarin fluorescence observed by d5 in *opt3-2* (**Fig. 1C, D**). In contrast, the coumarin-deficient *f6’h1-1* mutant responded to Fe deficiency under elevated pH with a strong induction of FCR activity, likely compensating for the lack of functional coumarin-mediated Fe mobilization (**Fig. 1C**). These trends emphasize that external pH, systemic Fe signals, and local Fe status jointly shape the balance and kinetics of FCR activity and coumarin biosynthesis in regulating Fe homeostasis.

Because *opt3-2* mutants exhibited a fluorescent coumarin halo under navFe (**Fig. 1D**) which is strikingly similar to previous reports in Arabidopsis (**Schmid et al., 2014; Tsai et al., 2018; Gautam et al., 2021**), the functional competence of these exuded coumarins was tested in a co-cultivation assay with *f6’h1-1* (**Fig. 1E**). When *f6’h1-1* seedlings were germinated and grown on navFe media between Col-0 and *opt3-2* seedlings, shoot greening and increased chlorophyll content were observed preferentially on the side facing *opt3-2*, indicating that coumarins secreted by *opt3-2* are more potent at mobilizing Fe than those released by Col-0 and are sufficient to partially rescue the Fe-deficiency symptoms in the coumarin-deficient background (**Fig. 1E, F**). The coumarin profile of Arabidopsis is known to include both fluorescent and non-fluorescent coumarins, whose relative contributions depend on external pH, Fe status, and genotypes under study (**Schmid et al., 2014; Tsai et al., 2018; Rajniak et al., 2018; Voges et al., 2019; Gautam et al., 2021; Robe et al., 2021; Paffrath et al., 2024)**. Fraxetin, scopoline, and scopoletin predominate under alkaline conditions, whereas sideretin is particularly important under acidic Fe deficiency (**Tsai et al., 2018; Rajniak et al., 2018; Gautam et al. 2021; Paffrath et al., 2024**). Transcriptomic analyses of the *opt3-2* mutant reveal extensive overlaps in the regulation of core Fe-responsive genes with that observed in transgenic lines overexpressing IRON MAN (IMA) peptides, which are known to regulate Fe uptake and homeostasis (**Grillet et al., 2018, 2023**). Notably, this shared regulatory signature includes key genes of the coumarin biosynthetic pathway, such as *MYB72, CYP82C4*, and *S8H* (**Grillet et al., 2023; Chia et al., 2023**). *IMAox* lines accumulate Fe (**Gautam et al., 2021; Grillet et al., 2018; Hirayama et al., 2018**), a phenotype closely resembling that of *opt3* mutants but with a difference in distribution of Fe in later (**Mendoza-Cózatl et al., 2014; Chia et al., 2023**). Consistent with this similarity, *opt3-2*, like the *IMAox* lines, can rescue the *f6’h1-1* mutant when grown in proximity on the navFe nutrient medium (**Fig. 1E, F**) (**Gautam et al., 2021**). Moreover, the UV-induced fluorescence pattern (365 nm) observed in *IMAox* lines grown under navFe conditions shows a striking resemblance to that of *opt3-2*, further supporting convergence in their Fe-deficiency responses (**Gautam et al., 2021**) (**Fig. 1D**). Together, these findings suggest that under navFe conditions, disruption of systemic Fe signaling in *opt3-2* may regulate Fe-responsive gene networks in a manner that is overlapping yet mechanistically distinct from IMA-mediated signaling, an idea that merits direct experimental testing.

Although this study focuses on three simple proxies, chlorophyll, FCR activity, and bulk coumarin fluorescence, the results point to substantial reprogramming of coumarin pathway flux and regulation in *opt3-2*, raising critical open questions about the underlying transcriptional control, the precise composition and abundance of individual coumarins, and their integration with OPT3-dependent systemic signaling. Overall, the work reveals a deeply intertwined crosstalk between reductive Fe uptake and coumarin-mediated Fe mobilization in Arabidopsis and demonstrates that disruption of systemic Fe signaling profoundly alters coumarin biosynthesis, its temporal deployment, and its functional impact on Fe nutrition in both the producing plant and neighboring roots.

## Material and Methods

### Plant materials and growth conditions

Seeds of *Arabidopsis thaliana* (L.) Heynh. ecotype Columbia-0 (Col-0) and the T-DNA insertion line *f6′h1-1* (SALK_132418C) were sourced from the Arabidopsis Biological Resource Center (ABRC, Ohio State University, USA). The *opt3-2* mutant (SALK_021168C) was kindly provided by Prof. David Mendoza-Cózatl (University of Missouri, USA). For surface sterilization, seeds were incubated for 5 min on a rocker in a solution containing 30% (v/v) commercial bleach (6% NaClO) and 70% (v/v) ethanol, followed by five washes with sterile Milli-Q water. Sterilized seeds were stratified for 2 d at 4 °C in darkness, sown on nutrient agar plates, and transferred to a controlled-environment growth chamber maintained at 21 °C under continuous illumination (50 μmol m^−2^ s^−1^).

Plants were grown on solid Estelle and Somerville (ES) medium (**Estelle and Somerville, 1987**) supplemented with 1.5% (w/v) sucrose and buffered with 1% (w/v) MES and solidified with 0.4% (w/v) Gelrite Pure (Kelco), as previously described by **Gautam et al., (2021**). Control (ES) medium contained 40 μM Fe-EDTA. Fe-deficient (-Fe) medium was prepared by omitting any Fe source and supplementing with 120 μM ferrozine to chelate trace Fe contaminants; the pH was adjusted to 5.5. For non-available Fe (navFe) conditions, Fe was supplied as 40 μM FeCl_3_ and the medium pH was adjusted to 7.0 using KOH. In navFe medium, MES was replaced with MOPS (1 g L^−1^) as the buffering agent.

### Chlorophyll content and ferric chelate reduction activity

Freshly harvested shoots were ground in liquid nitrogen, and chlorophyll was extracted with 80% acetone. Total chlorophyll (from absorbance at 645 and 663 nm) was estimated using a protocol from **Wellburn (1994)**.

Ferric chelate reductase (FCR) activity was measured for pooled roots from 5-10 intact seedlings as per the protocol described by **Grillet et al., (2014)**. Roots were incubated in the dark for 30 min in an assay solution containing 100 µM Fe-EDTA and 300 µM bathophenanthroline disulfonate (BPDS) dissolved in 10 mM MES buffer (pH 5.5). Following incubation, the formation of Fe^2+^-BPDS_3_ complex was quantified by measuring absorbance at 535 nm using a PowerWave XS2 microplate reader (BioTek Instruments). Fe^2+^ concentrations were calculated from a standard curve. All experiments were performed with at least three independent biological replicates.

### Detection of fluorescent compounds in roots and media

The accumulation of fluorescent compounds in roots and their secretion into the growth medium were visualized using a BioSpectrum 600 imaging system (UVP). Samples were imaged with an excitation wavelength of 365 nm and an emission filter range of 485-655 nm (SYBR Gold), with an exposure time of 9 s, as previously described (**Tsai et al., 2018**).

### Co-cultivation assay to rescue coumarin-deficient mutant

Rescue of the coumarin-deficient *f6′h1-1* mutant was assessed using a co-cultivation assay on navFe medium. *f6′h1-1* seedlings were germinated and grown in proximity to Col-0 and *opt3-2* seedlings for 14 d, arranged horizontally at equal distances from one another on the same navFe plate. The phenotype was observed at 14d, and the shoot was harvested to quantify chlorophyll.

### Statistical analyses

All statistical analyses were performed in R version 4.0.3 (**R Core Team, 2020**). Data reshaping was carried out using the reshape2 package (v1.4.4; **Wickham, 2007**). One- and two-way analyses of variance (ANOVA) were conducted using the stats package (**R Core Team, 2020**), and multiple comparisons were performed using the multcomp package (v1.4-16; **Hothorn et al., 2008**). Post-hoc comparisons were conducted using Tukey’s HSD test (TukeyHSD, stats), and compact letter displays were generated using the glht function (multcomp). Graphs were generated using GraphPad Prism version 9.

## Acknowledgements

The author would like to thank Dr. Wolfgang Schmidt (a retired Research Fellow at the Academia Sinica, Taiwan) for inspiring the initial research questions and supporting this work through a grant. We thank Prof. David Mendoza-Cózatl (University of Missouri) for providing seeds of the *opt3-2* mutant.

## Statements and Declarations

### Ethical Approval

Not applicable.

